# Enhanced C/EBPs binding to C>T mismatches facilitates fixation of CpG mutations

**DOI:** 10.1101/2020.06.11.146175

**Authors:** Anna S. Ershova, Irina A. Eliseeva, Oleg S. Nikonov, Alla D. Fedorova, Ilya E. Vorontsov, Dmitry Papatsenko, Ivan V. Kulakovskiy

**Author notes:** co-first authors, equal contribution (Anna S. Ershova, Irina A. Eliseeva). deceased (Dmitry Papatsenko). E-mails Ivan V. Kulakovskiy -, Dmitry Papatsenko - deceased, Ilya E. Vorontsov -, Alla D. Fedorova -, Oleg S. Nikonov -, Irina A. Eliseeva -, Anna S. Ershova.

## Abstract

Knowledge of mechanisms responsible for mutagenesis of adult stem cells is crucial to track genomic alterations that may affect cell renovation and provoke malignant cell transformation. Mutations in regulatory regions are widely studied nowadays, though mostly in cancer. In this study, we decomposed the mutation signature of adult stem cells, mapped the corresponding mutations into transcription factor binding regions, and assessed mutation frequency in sequence motif occurrences. We found binding sites of C/EBP transcription factors strongly enriched with [C>T]G mutations within the core CG dinucleotide related to deamination of the methylated cytosine. This effect was also exhibited in related cancer samples. Structural modeling predicted enhanced CEBPB binding to the consensus sequence with the [C>T]G mismatch, which was then confirmed in the direct experiment. We propose that it is the enhanced binding of C/EBPs that shields C>T transitions from DNA repair and leads to selective accumulation of the [C>T]G mutations within binding sites.

## Introduction

Accumulation of somatic mutations leads to cancer and other diseases (Blokzijl et al., 2016). Different organs and tissues exhibit different probability to develop cancer, which can be explained by the number of divisions of the respective adult stem cell (ASC) (Tomasetti and Vogelstein, 2015). Thus, studying of mutational processes in stem cells is crucial to understand the tumorigenesis (Blokzijl et al., 2016; Franco et al., 2019; Rouhani et al., 2016; Saini and Gordenin, 2018; Yoshihara et al., 2017).

Distribution of somatic mutations across a genome varies depending on a mutation class and underlying mutational processes, chromatin organization (Schuster-Böckler and Lehner, 2012), DNA replication timing (Stamatoyannopoulos et al., 2009; Woo and Li, 2012), and activity of repair systems (Supek and Lehner, 2015). Point mutations are depleted within transcription factor binding sites in induced pluripotent stem cells (iPSCs) (Yoshihara et al., 2017) and in certain cancers (Rheinbay et al., 2020; Vorontsov et al., 2016; Rheinbay et al., 2017). Nevertheless, alterations in regulatory regions are associated with many complex traits (Deplancke et al., 2016). Particularly, there are recurrent functional mutations in regulatory regions (Saini and Gordenin, 2018), such as the well-studied recurrent C>T transition that creates a strong binding site for the GABP transcription factor in the TERT promoter and is associated with increased cancer risk (Bell et al., 2015; Horn et al., 2013; Vinothkumar et al., 2020).

Somatic mutations are caused by a combination of DNA damage and DNA repair failures (Volkova et al., 2020). The interplay of these processes can produce mutation sets in different preferred genomic contexts, the so-called mutation signatures (Alexandrov et al., 2020, 2013), with varying impact on creation or disruption of transcription factor binding motifs (Yiu Chan et al., 2019). In cancer, the overall patterns of mutational processes in transcription factor binding sites are very complex, due to the interference of numerous incidental circumstances, both local, such as the extended context of mutation signatures (Fredriksson et al., 2017), and global, such as the pressure of clonal selection (Vorontsov et al., 2016).

Data on mutations in genomes of human adult stem cells (hASCs) from the healthy donors (Blokzijl et al., 2016) provide a unique opportunity to study a mutation process in the absence of major selection pressure and thus focus on molecular mechanisms targeting particular genomic sites. In this study, by analyzing mutations within transcription factor binding sites in hASCs (Blokzijl et al., 2016) and iPSCs (Rouhani et al., 2016), we identified the binding sites of the C/EBP transcription factors as mutation hotspots. Experimental verification of altered C/EBP binding in case of the C>T mismatch (T-G pairing) and the mutated base pair (canonical T-A, post replication) in the CpG context suggests a molecular mechanism of targeted mutagenesis acting in hASCs and cancer cells.

## Results and Discussion

### Transcription factor binding regions encompass up to a third of mutations in human adult stem cells

To distinguish features of different mutational processes, we decomposed the stem cell mutations into three distinct signatures (Fig EV1A) (Alexandrov et al., 2013; Blokzijl et al., 2016).

Signature 1 (called SBS1 in (Alexandrov et al., 2020) and Sig.B in (Blokzijl et al., 2016)) represents mainly the C to T transitions in CpG contexts resulting from spontaneous deamination of methylated cytosine into thymine (Alexandrov et al., 2013; Blokzijl et al., 2016). This mutational process plays a major role in small intestinal and colon ASCs (Fig EV1B).

Signature 2 (C>A, similar to signature SBS18 in (Alexandrov et al., 2020) and Sig.C in (Blokzijl et al., 2016)) is related to reactive oxygen species. As a result of oxidative DNA lesions, guanine changes to 8-oxoguanine that can mispair with adenine and create the G:C > T:A transversions. This signature was associated with mutations in reparation protein MUTYH in colorectal cancers (Viel et al., 2017) and often detected in cell cultures *in vitro* (Phillips, 2018).

Signature 3 (corresponding to signature SBS5 in (Alexandrov et al., 2020) and Sig.A in (Blokzijl et al., 2016)) is characterized by T:A to C:G transitions and mainly found in liver samples (see EV1B). The mechanism of this signature is unknown, Blokzijl and colleagues suggested this signature to be associated with aging (Blokzijl et al., 2016).

Based on the trinucleotide sequence context, each point mutation was assigned to the signature where its contribution was the highest. Next, we used the map of human transcription factor binding regions (the cistrome) (Vorontsov et al., 2018) to estimate the fraction of mutations falling into regions potentially bound by transcription factors. For each signature, we found about 30% of mutations within cistrome regions (Table EV1), which allowed us to perform a detailed analysis of mutations within the occurrences of particular transcription factor binding motifs, i.e. the predicted transcription factor binding sites.

### Mutations in C/EBP binding regions are precisely enriched within binding sites

To analyze the preferred locations of mutations relative to occurrences of transcription factor binding motifs, we performed a comprehensive scan of [−100;+100] bp windows anchored at the mutated bases with the motif models from the HOCOMOCO v11 database (Kulakovskiy et al., 2018). The motif occurrence with the highest score was marked in each window, and only windows with those best hits passing a P-value of 0.0005 were taken for further analysis. The anchoring mutations and the respective windows were classified using two binary features: whether located within/outside of the cistrome (Vorontsov et al., 2018) and carrying the mutation within/outside of the motif occurrence. This allowed estimating relative enrichment or depletion of mutation within and outside of the motif occurrences depending on the location of the tested window (within or outside of the cistrome) with Fisher’s exact test. We performed this analysis separately for three mutation signatures for each sample. Despite statistical power limited by low mutation frequency in ASCs, we found two significant effects, both arising from the S1 signature (Table EV2). First, there were a number of Zinc-finger proteins, mostly from SP- and KLF-families binding CCCCG-boxes. Those motif occurrences, when found within the cistrome, were devoid of mutations, suggesting that direct binding of these proteins could play a protective role, e.g. by reducing methylation (Long et al., 2016). Second, we found the C/EBP and C/EBP-related motif repeated the analysis using the CEBP-only subset of the cistrome and the subset of [C>T]G-only mutations as the primary component of the S1 signature. In this setting, the effect was even better exhibited, with the mutation rate within the C/EBP motif occurrences being 3 to 5 times higher than expected (Fig 1A). Next, we analyzed the positional preferences of mutations in the vicinity of the motif occurrences and found a strongly increased mutation rate in the core CG within the occurrences of the C/EBP binding motif (Fig 1B).

**Figure 1.**
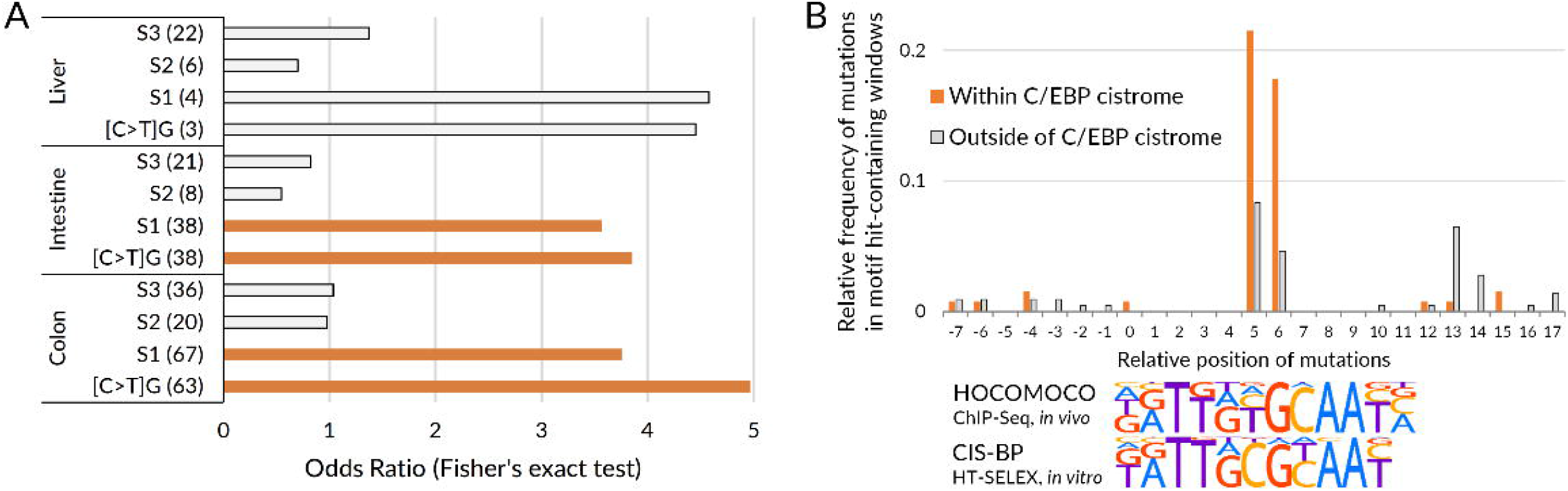
C/EBP binding sites are enriched with [C>T]G somatic mutations in human adult stem cells. **A** Odds ratios (X-axis) demonstrating the enrichment of hASC mutations within C/EBP motif occurrences in the cistrome regions. Bar groups denote different hASC samples (liver, intestine, and colon). Bars within each group correspond to particular mutation groups (signatures S1-3, and [C>T]G subset), the number of mutations within C/EBP occurrences in the cistrome is shown in brackets. Orange bars correspond to the P-values of less than 0.05 (Fisher’s exact test). **B** Frequency of mutations (Y-axis) relative to the C/EBP motif occurrences (X-axis). Only [C>T]G mutations are considered. Logos depict optimal C/EBP motifs recognizing *in vivo* (HOCOMOCO) and *in vitro* (CIS-BP) sites.

C/EBPs are known to prefer m^5^C within the core CG pair (Sayeed et al., 2015). Therefore, their functional binding sites can be associated with the elevated CpG-methylation-dependent mutagenesis. Yet, considering only S1 or its [C>T]G subset, we found enrichment of mutations within the ‘genuine’ C/EBP binding sites (the motif occurrences within the cistrome) against the non-cistrome control. With this setup, the observed effect cannot be attributed solely to high ‘background’ mutability of methylated CpGs (since we compare mutations of the same signature within and outside of motif occurrences) or to some unknown extended context preferred by mutations (since we compare occurrences of the same sequence motif within and outside of the cistrome). Thus, the enriched mutation within C/EBP binding sites is directly associated with the C/ EBP DNA binding.

Analysis of mutations in the related cancer samples (partly sharing the mutation signatures of the analyzed stem cells, Fig EV2A) fully confirmed the targeted mutagenesis of CpGs within C/EBP motif occurrences (Fig EV2B). Previously, a similar effect was reported for breast cancer data in (Melton et al., 2015), suggesting that it is a common characteristic feature of [C>T]G mutations in regulatory regions.

There are two possible explanations for the enrichment of ASC and tumor mutations within core CG pairs of the C/EBP binding sites. First, the C/EBP binding could directly facilitate cytosine deamination within methylated CpGs. However, it seems unlikely for a transcription factor to have such a direct mutagenic activity. Second, the DNA-bound protein could be playing a protective role by making the respective DNA region inaccessible to other cellular machinery, i.e. tight C/EBP binding could protect a spontaneously mutated site from the mismatch repair. This scenario could be realized through an increased protein affinity to certain mismatches, which was reported for many transcription factors (Afek et al., 2019). C/EBP-related proteins were not explored in this regard, and we hypothesized that it is the enhanced C/EBP binding that protects the sites with mismatches from the co-transcriptional repair and allows for fixation of the respective mutations at the replication stage.

### Structural analysis predicts increased CEBPB affinity to single-strand [C>T]G mismatches

To explore CEBPB binding to [C>T]G mismatches, we performed structural modeling. We constructed two models of CEBPB-DNA complexes with a consensus nucleotide sequence (Fig 2A). In the first model, the cytosine in the CG pair was replaced with thymine in one of the chains of the DNA duplex. In the other model, we additionally replaced the complementary guanine with adenine, thus restoring the canonical base pairing.

**Figure 2.**
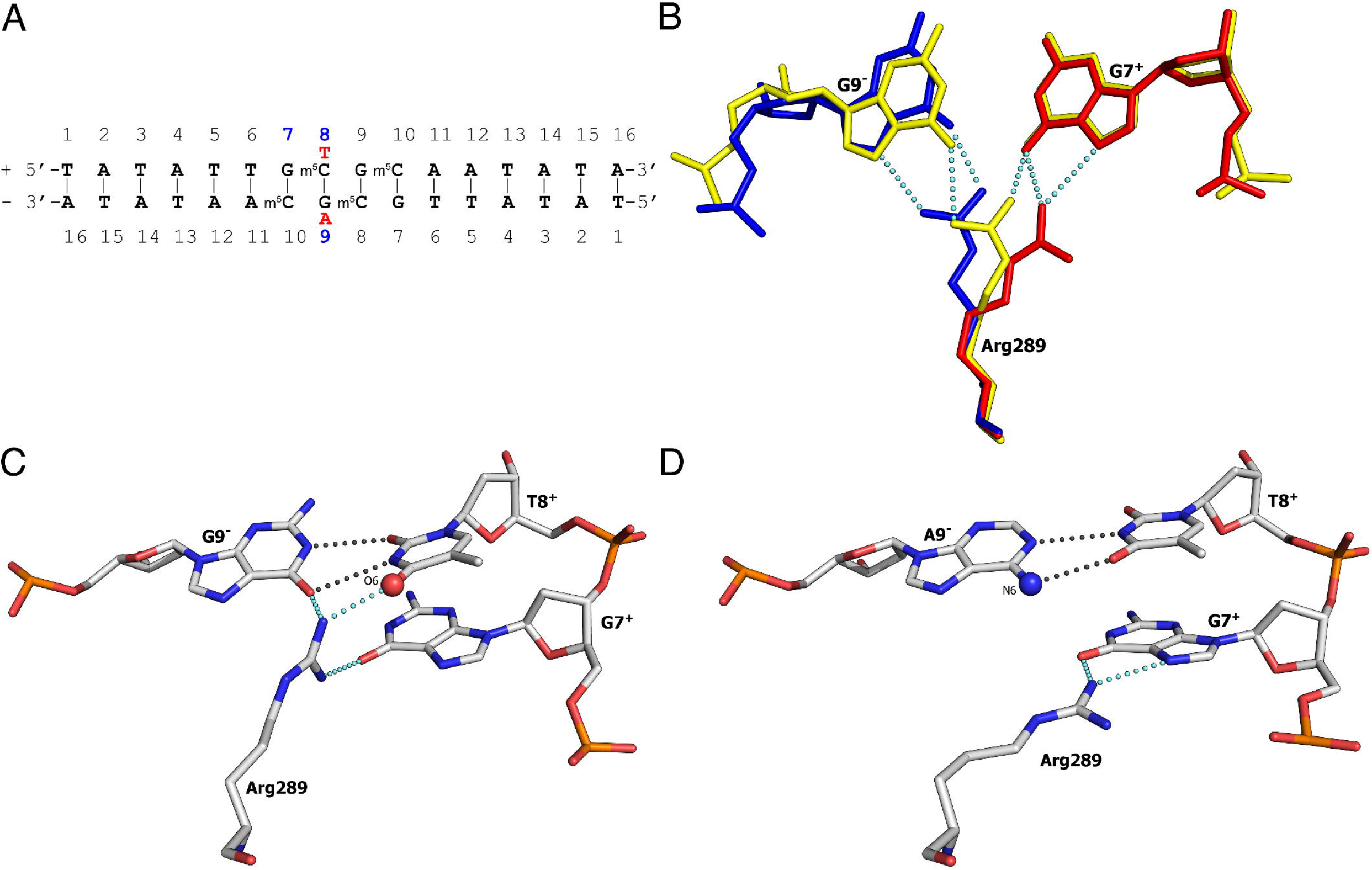
Additional hydrogen bond between Arg289 and O6 of the introduced thymine and GT wobble pairing predict enhanced CEBPB binding to [C>T]G mismatches. **A** DNA duplex used for modeling of CEBPB-DNA complexes. Positions of nucleotides interacting with Arg289 are marked in blue, nucleotide substitutions are shown in red, (+) and (–) denote DNA chains. **B-D** Interactions of CEBPB Arg289 with DNA. Hydrogen bonds are shown by blue dotted lines (Arg289-DNA interactions) and gray dotted lines (nucleotides base pairing). (**B**) Original PDB structures: 6mg1 (blue), 6mg2 (yellow), 6mg3 (red). Molecular modelling: (**C**) C8^+^>T8^+^ substitution, the O4 atom of T8^+^ is shown as the red sphere and labeled; (**D**) C8^+^>T8^+^ and G9^−^>A9^−^ substitutions, the N6 atom of A9^−^ is shown as the blue sphere and labeled.

Three structures of the CEBPB-DNA complexes defined at atomic resolution (PDB codes: 6mg1, 6mg2, 6mg3 (Yang et al., 2019)) were taken as the basis for modeling. In these structures, the double-stranded DNA has the palindromic sequence TATATTGCGCAATATA, i.e., the core consensus sequence TTGCGCAA occupies positions from 5 to 12. The cytosines have a methyl group attached at position 5.

We arbitrarily named one of the chains of the DNA duplex as the (+) chain, and the other as (−) chain, so T1^+^ denoted the first nucleotide of the (+) chain, and T1^−^ denoted the first nucleotide of the (−) chain (Fig 2A). Under this notation, the C>T and G>A substitutions in the core consensus are C8^+^>T8^+^ and G9^−^>A9^−^ (Fig 2A). Models with the single-strand C8^+^>T8^+^ and double-strand (C8^+^>T8^+^ and G9^−^>A9^−^) substitutions were obtained without noticeable changes in the overall geometry of the DNA duplex.

Analysis of the initial structures shows that Arg289, which plays a key role in specific CEBPB– DNA binding (Yang et al., 2019), can interact with the nucleic acid in different ways. The main binding site for Arg289 is formed by the second guanine of the consensus sequence (G9^−^). Arg289 can interact with G9^−^ in two ways: it can either form hydrogen bonds with the N7 and O6 atoms or bind to the O6 atom of G9^−^ and the O6 atom of the G7^+^, occupying an intermediate position between G9^−^ and G7^+^ (Fig 2B). The interactions of Arg289 with G9^−^ are stabilized by the Van Der Waals interactions of the Arg289 side chain with the Val285 side chain and the methyl group of the methylated C8^−^ (m^5^C8^−^) (Yang et al., 2019). When these interactions are weakened (e.g., through replacement of Val285 by Ala285), Arg289 can completely exchange hydrogen bonds with G9^−^ for hydrogen bonds with G7^+^, more precisely, with the N7 and O6 atoms of G7^+^ (Fig 2B). There are no steric restrictions for such interaction even if the above mentioned Van Der Waals contacts are preserved. However, the hydrogen bonds with G9^−^ are accessible to the solvent to a much lesser extent than the hydrogen bonds with G7^+^, which makes the contact with G9^−^ more stable. Thus, Arg289 can switch between the acceptor groups of G9^−^, G7^+,^ and water due to the accessibility of the hydrogen bonds formed with DNA to the solvent, but the position of Arg289 is more stable in case of the interaction with G9^−^.

The molecular modeling shows that the C8^+^>T8^+^ substitution and the formation of a Wobble G9^−^-T8^+^ pair (Ho et al., 1985) in the contact area of Arg289 with G9^−^ and G7^+^ introduces an additional strong acceptor of the hydrogen bond, namely the O4 atom of T8^+^. This acceptor insertion increases the chances that during the exchange of acceptor groups, Arg289 will retain DNA as a partner for the formation of hydrogen bonds and stabilizes its side chain in a position convenient for interaction with G9^−^ (Fig 2C). Thus, the DNA segment with the [C>T]G mismatch should exhibit stronger affinity to CEBPB.

An additional substitution in the (−) chain (G9^−^ >A9^−^) leads to the formation of the canonical A9^−^-T8^+^ pair. In this case, the acceptor of the hydrogen bond (atom O6 of G9^−^) for Arg289 is replaced by the hydrogen donor group NH2 (N6 atom of A9^−^) (Fig 2D). This substitution prevents the formation of hydrogen bonds with Arg289 in 9^−^ position and reduces the chances of Arg289 to retain DNA as a partner for the hydrogen bonds formation. The formation of the canonical A9^−^-T8^+^ pair causes Arg289 interaction with G7^+^ and formation of hydrogen bonds that are more accessible to the solvent than in the case of interactions with G9^−^. Thus, the binding site with a double-strand substitution should have significantly lower affinity to CEBPB.

### CEBPB has a strong affinity to consensus sites with single-strand [C>T]G DNA mismatches

To verify bioinformatics prediction, we performed an EMSA experiment with nuclear extract from HEK293T cells expressing FLAG-CEBPB (LAP2 isoform, Fig 3A) and synthetic oligonucleotides containing a palindromic consensus or mutated C/EBP binding site (Fig 3B). CEBPB was selected as a representative member of the C/EBP family since its motifs demonstrated the best family-wide recognition both *in vivo* and *in vitro* (Ambrosini et al., 2020). We used non-methylated oligonucleotides, as well as oligonucleotides carrying m^5^C within the core CG pair. To test the CEBPB binding, we used radiolabeled oligonucleotides with the methylated consensus CEBPB binding site (wt_m^5^C); other oligos were used as unlabeled competitors. The presence of CEBPB was confirmed by the formation of a low-motility DNA-protein complex with anti-FLAG antibodies (Fig 3C).

**Figure 3.**
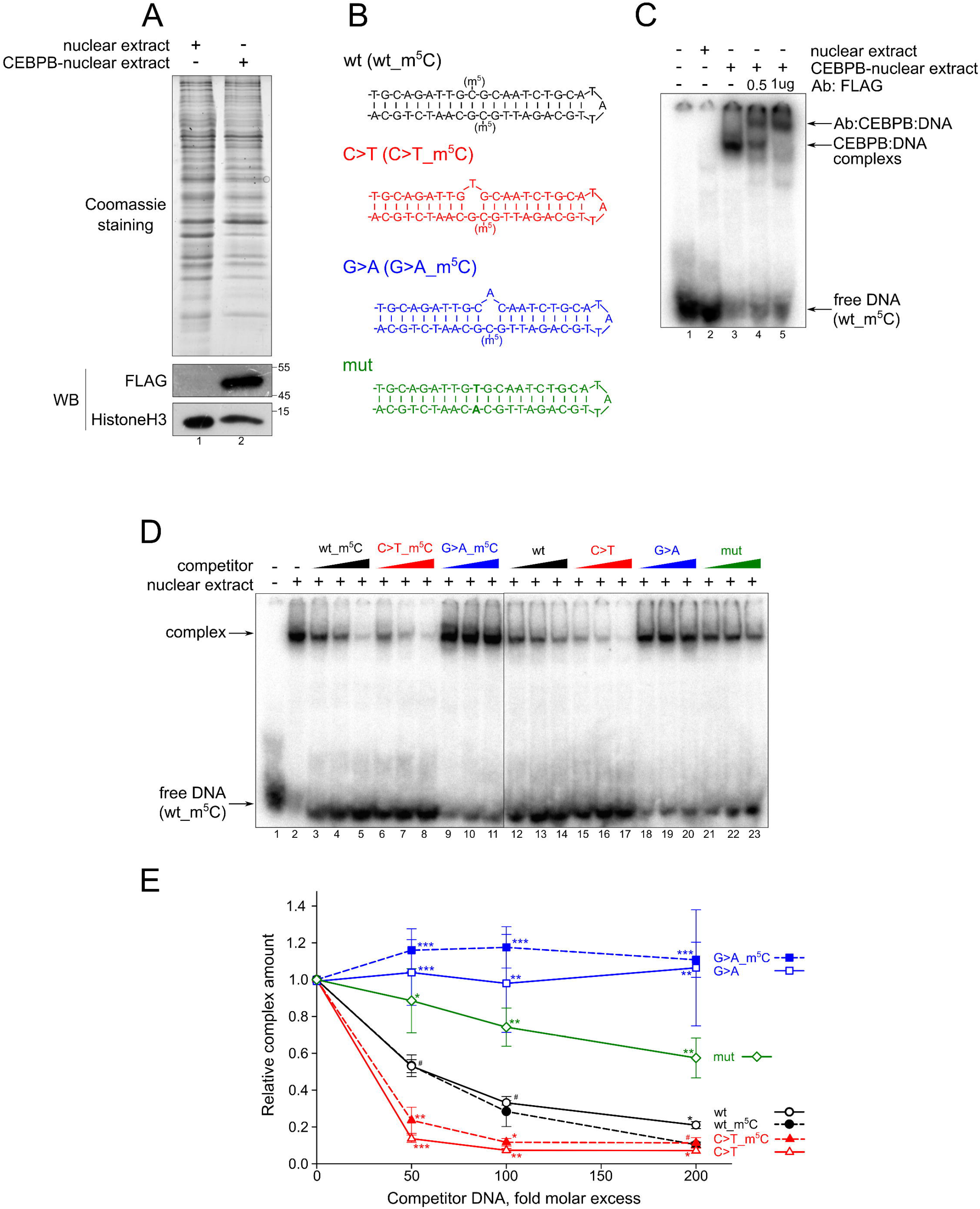
CEBPB has increased affinity to the [C>T]G mismatches and low affinity to the respective double-strand substitutions. **A** Nuclear extracts from HEK293T (line#1) or HEK293T expressed FLAG-LAP2 (an isoform of CEBPB, line#2) were separated by SDS-PAGE and stained with Coomassie Blue or subjected to Western blotting using antibodies against FLAG (anti-DDDDK) and Histone H3 (loading control). **B** Oligonucleotides used in EMSA experiments. The position of m^5^C and the oligonucleotide names are presented in brackets. **C** Nuclear extracts from HEK293T or HEK293T expressed FLAG-LAP2 (CEBPB) were pre-treated for 20 min at 30°C with nonspecific competitor DNA in the absence or presence of anti-FLAG antibodies. To form DNA-protein complexes radiolabeled wt_m^5^C oligonucleotide was incubated with or without pretreated nuclear extracts for 20 min at 30°C. The DNA-protein complexes were separated in native 4% PAAG and visualized by autoradiography. **D** Nuclear extract from HEK293T expressed FLAG-LAP2 (CEBPB) was pretreated with non-specific competitor DNA for 20 min at 30°C. To form DNA-protein complexes radiolabeled wt_m^5^C oligonucleotide was incubated with or without pretreated nuclear extracts for 20 min at 30°C in the absence or presence of unlabeled competitor oligonucleotides (at 50-, 100- or 200-fold molar excess). The DNA-protein complexes were separated in native 4% PAAG and visualized by autoradiography. The relative amount of radioactivity was determined using a Packard cyclone Storage phosphor System. **E** The quantification results. The radioactivity of DNA-CEBPB complexes was normalized to the radioactivity of the complex without competitor oligonucleotides. Values are the mean of at least three independent experiments. Two-tailed Student’s t-test was used to estimate the statistical significance of the difference in the relative complex amount formed in the presence of a particular competitor versus the wt_m^5^C competitor. ***p<0.001, **p<0.001, *p<0.05.

As shown in Figs 3D-E, oligonucleotides with the single-strand G>A mismatch and double-strand C>T(G>A) point mutation (mut) have a significantly lower affinity to CEBPB and weakly compete for its binding. An effect of m^5^C methylation on the CEBPB binding is rather minor, although the respective oligonucleotides act as slightly better competitors compared to the same non-methylated sequences, in agreement with (Sayeed et al., 2015). In contrast, oligos with the single-strand C>T mismatch compete for the CEBPB binding much stronger than the canonical ‘wild type’ palindromic CG-carrying oligos.

### Model of selective fixation of [C>T]G mutations through enhanced C/EBP binding

We discovered elevated mutagenesis of CpGs within C/EBP binding sites in hASCs. Because the studied sets of mutations in ASCs reflect processes active in cells of healthy donors, thus the effect cannot arise from positive selection. The purifying selection could lead to the depletion but not enrichment of mutations in particular positions of the sites. Thus, there should be a molecular mechanism mutating C/EBP sites in normal hASCs and in cancer cells similarly. We propose that it is the enhanced C/EBP binding that shields single-nucleotide [C>T]G mismatches from co-transcriptional repair so mutation fixation becomes possible at the replication stage (Fig 4).

**Figure 4.**
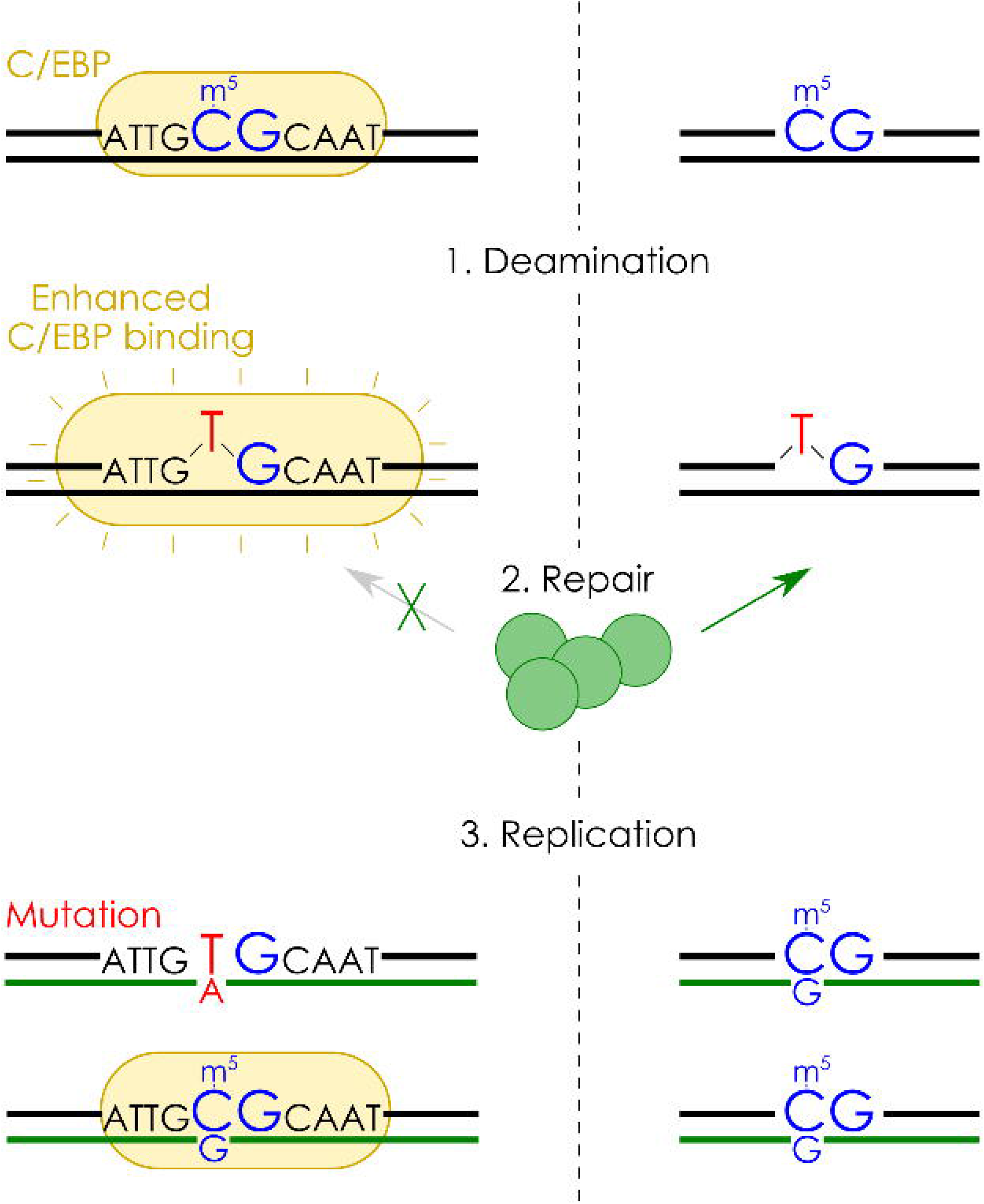
The proposed model of [C>T]G mutation fixation through enhanced C/EBP binding to single-nucleotide mismatches.

Among multiple motifs, in the cistrome analysis (particularly, in Fig 1B), we used the CEBPB motif (ID: CEBPB_HUMAN.H11MO.0.A) of the HOCOMOCO database (Kulakovskiy et al., 2018), which was the best C/EBP-family motif in terms of ChIP-Seq peaks recognition (Ambrosini et al., 2020). However, an alternative motif from CIS-BP (ID: M05840_2.00) (Weirauch et al., 2014) was found the best in recognizing the binding sites from *in vitro* data. When position along each other in the plot (Fig 1B), the major CG dinucleotide in the palindromic *in vitro* consensus (the motif from CIS-BP) corresponded to TG *in vivo* (the motif from HOCOMOCO). This is consistent with the case of considering not only the single best but also multiple ChIP-Seq (*in vivo*) and HT-SELEX (*in vitro*) C/EBP motifs presented in CIS-BP database (Weirauch et al., 2014), suggesting that many TG-carrying weaker sites originated from canonical CG-carrying ones by point mutations.

Yet, the genomic dinucleotide composition is devoid of CGs (CG to TG ratio of 1:7), and the core of the C/EBP consensus motif *in vivo* is relatively enriched with CGs (CG to TG ratio of 1:2, see the respective position count matrix in HOCOMOCO). These general data agree with promoter-level estimates for motif subtypes in cistrome-overlapping and non-overlapping promoters. Particularly, the promoters overlapping CEBP-cistrome exhibited 7.1 times higher CG-to-TG ratio (CG:TG=1133:2588), comparing to the remaining set of promoters (CG:TG=96:1563) when examining the best occurrences of the best C/EBP binding motif based on *in vivo* ChIP-Seq data (HOCOMOCO CEBPB_HUMAN.H11MO.0.A). The same effect, although of a lower magnitude (2.7 times more CGs, CG:TG=2006:1242 versus CG:TG=340:565), was found when using the best *in vitro* motif (CIS-BP M05840_2.00). Thus, in the reference human genome the canonical CG-containing C/EBP sites in the C/EBP cistrome appear specifically conserved. Probably, they avoid mutations either as being rarely methylated (e.g. located in hypomethylated CpG-islands), or through global purifying selection at the population level.

To evaluate if the CG- and TG-carrying subtypes are functionally distinct, we classified promoters according to the subtypes of the best C/EBP motif occurrences and performed the pathway enrichment analysis (see Methods). We found that the genes with the CG-subtype C/EBP sites in promoters are consistently associated with the ‘RNA metabolism’ (Reactome R-HSA-8953854), showing log10(adjusted P-value) from 4.3 to 11.6 for different combinations of the cistrome subset (CEBPA/B) and motif (*in vivo*/*in vitro*). This association was never found among top significant terms for TG-subtype C/EBP sites. The same CG- but not TG- association (although with lower significance) was found for ‘ribonucleoprotein complex biogenesis’ (GO:0022613).

Current data on the C/EBP proteins, including CEBPA and CEBPB, suggest that they are tightly involved in establishing the methylation status of the regulatory regions (Schäfer et al., 2018), which is linked to high-level processes such as energy metabolism and longevity (Niehrs and Calkhoven, 2020). Targeted somatic mutagenesis of C/EBP sites in adult stem cells might be yet another contribution to aging or malignant cell transformation.

The widest repertoire of mutation signatures can be found in cancer cells. Enhanced binding of transcription factors possibly allows for fixation of mutations from various signatures, thus canalizing mutation-induced changes of regulatory networks in transcription factor-dependent and signature-specific mode. Particularly, strongly bound mismatches may provide a brief window of opportunity (post-mismatch but pre-replication) for a single cell to ensure significant down- or up-regulation of a particular gene, if the mismatch occurs in the binding site within a critical regulatory region such as the core promoter. A distinct functional outcome of a DNA mismatch has the potential to drive the clonal evolution of cancer cells or ASC cell transformation, thus motivating further analysis of regulatory genomic alterations in other types of hASCs and cancers.

## Materials and Methods

### Bioinformatics analysis

#### Overview of mutation data sets

We used the mutations data for hASC (Blokzijl et al., 2016), iPSC (Bhutani et al., 2016; Rouhani et al., 2016), and related cancer samples (mutation calls from the whole-genome sequencing experiments were downloaded from the ICGC data portal (Zhang et al., 2019)). For cancer samples, the recurrent mutations were merged. An overview of the data is presented in Table EV1.

#### Analysis of mutation signatures

The occurrences of all 96 trinucleotide contexts were counted for each dataset using the Mutational Patterns R/Bioconductor package (Blokzijl et al., 2018). Then, using the same package, the mutational signatures S1-S3 for hASC and iPSC samples were extracted from 96 trinucleotide contexts by non-negative matrix factorization (NMF) with the ‘extract_signatures’. A possible impact from each signature on mutational profiles of cancer samples was then estimated with cosine similarity ‘cos_sim’.

For detailed analysis, each particular mutation was assigned to a single signature based on the trinucleotide context of the mutation. We performed this step for contexts whose relative contribution to the selected signature was more than 5% higher than their average relative contribution to all signatures. Because two contexts (A[C>A]C and A[T>A]T) did not pass the thresholds, they were not assigned to particular signatures, and the respective mutations (~2% of total) were omitted from the downstream analysis.

#### Analysis of mutations in transcription factor binding sites

Human hg19 cistrome data (genomic regions of transcription factor binding) for 599 human transcription factors (TFs) (Vorontsov et al., 2018) were used for the mutation enrichment analysis. The complete cistrome was constructed from high reliability cistromes (A, B, C, see (Vorontsov et al., 2018) for details) of all TFs through merging with bedtools v2.27.1 (Quinlan and Hall, 2010). C/EBPs-only cistrome was obtained in the same manner considering only C/EBPA, C/EBPB, C/EBPD, C/EBPE, and C/EBPG binding regions.

The sequence motif analysis was performed with 402 position weight matrices from the HOCOMOCO v11 HUMAN CORE database (Kulakovskiy et al., 2018). Motif finding was performed in 101 bp windows centered at mutations with the SPRY-SARUS software. The motif occurrence thresholds were selected according to motif P-value of 0.0005 as in (Vorontsov et al., 2018), roughly resulting in 1 expected random hit per ten 101 bp windows. An additional motif based on *in vitro* high-throughput SELEX data was downloaded from CIS-BP (Weirauch et al., 2014). Two-tailed Fisher’s exact test using 2×2 contingency tables was performed independently for each weight matrix, the resulting P-values were corrected for the number of multiple tested motifs using the Benjamini-Hochberg (FDR) procedure. The complete cistrome of all transcription factors was used in the initial analysis (Fig 1A), the C/EBPs-only cistrome was used for the detailed analysis (Figs 1B,C, Fig EV2).

#### Analysis of C/EBP motif subtypes in promoters

In this analysis, we considered protein-coding genes with the cistrome-overlapping promoters ([−400,+100]bp relative to the transcription start sites annotated in GENCODE v34 basic annotation (Frankish et al., 2019)). CEBPB_HUMAN.H11MO.0.A and M05840_2.00 motifs were analyzed separately. The best motif hit was selected in each promoter, only the hits passing P-value of 0.0005 were taken for further analysis. The most reliable CEBPA and CEBPB cistromes were used to annotate promoters and assemble gene sets for the gene enrichment analysis, which was performed with Metascape (Zhou et al., 2019) (default parameters).

#### Structural modeling

Three structures of the CEBPB-DNA complexes defined at atomic resolution (PDB codes: 6mg1, 6mg2, 6mg3 (Yang et al., 2019)) were used for modeling. The structural models of the DNA fragment mutant forms were manually built in the Coot software with the homologous modeling (Emsley et al., 2010). Local geometry changes upon nucleotide substitutions were corrected using the “regularize zone” function until idealized values of the angles and bond lengths were achieved.

### Experimental verification

#### Preparation of the nuclear extract

HEK293T cells (originally obtained from ATCC, American Type Culture Collection) were kindly provided by Dr. Elena Nadezhdina (Institute of Protein Research, Russian Academy of Sciences, Pushchino, Russia). The cells were cultivated by a standard method in DMEM (Dulbecco Modified Eagle Medium) supplemented with 10% fetal bovine serum, 2 mM glutamine, 100 U/mL penicillin, and 100 µg streptomycin (PanEco, Moscow, Russia). The cells were kept at 37°C in a humidified atmosphere containing 5% CO_2_.

To obtain CEBPB-expressing HEK293T, the cells were transfected with 12 µg per 10 sm dish of pCMV-FLAG LAP2 (the long isoform of CEBPB) using Lipofectamine 3000 (Thermo Fisher Scientific). After 24h the cells were plated from one 10 sm dish to three and cultivated under standard conditions for additional 48h. The pCMV-FLAG LAP2 plasmid was a gift from Joan Massague (Addgene plasmid #15738; http://n2t.net/addgene:15738; RRID:Addgene_15738) (Gomis et al., 2006).

To prepare nuclear extract, the cells were washed two times and collected in ice-cold PBS. Then the cells were lysed in five pellet volumes of buffer A (10 mM Hepes 7.6, 10 mM KCl, 1.5 mM MgCl_2_, 1 mM DTT). The lysates were passed ten times through a 26G needle, incubated on ice for 10 min, and centrifuged at 4°C at 16,000 g for 5 min. The nucleus pellet was washed two times with two volumes of buffer A. Each time, the lysates were passed tenfold through a 26G needle, incubated on ice for 10 min, and centrifuged for 5 min. The nucleus was lysed in two volumes of buffer C (20 mM Hepes 7.6, 420 mM NaCl, 1.5 mM MgCl_2_, 1 mM DTT, 20% glycerol) for 2h at 4°C with agitation. The nuclear extract was cleared by centrifugation at 4°C at 16,000 g for 15 min and stored at −80°C.

#### Western blot and antibodies

For the Western blot analysis, the nuclear extract was supplemented with SDS electrophoresis sample buffer, separated by SDS-PAGE, and stained with Coomassie Blue or transferred onto a nitrocellulose membrane. The membrane was blocked for 1h at room temperature with 5% nonfat milk in TBS-T (10 mM Tris-HCl, pH 7.6, 150 mM NaCl, 0.1% Tween 20) and incubated overnight at 4°C in TBS-T supplemented with BSA (5%) and appropriate antibodies. The membrane was then washed three times with TBS-T, incubated for 1 h with 5% nonfat milk in TBS-T and horseradish peroxidase-conjugated goat anti-rabbit IgG (1:4000, #7074, CST) and then washed three times with TBS-T. The immunocomplexes were detected using an ECL Prime kit (GE Healthcare) according to the manufacturer’s recommendations. The rabbit primary antibodies anti-DDDDK tag (binds to FLAG, 1:2500, #ab1162, Abcam) and anti-histone H3 (1:10000, #4499S, CST) were used in the process.

#### Electrophoretic mobility shift assay (EMSA)

Methylated (m^5^C) oligonucleotide with consensus palindromic C/EBP binding motif (wt_m^5^C, Table 1) was 5’-radiolabeled with T4-polynucleotide kinase in the presence of [γ-^32^P]-ATP (4,000 Ci/mM; IBCh, Russia) according to the manufacturer’s recommendations. The oligonucleotide was purified by gel filtration using Illustra ProbeQuant G-50 Micro Columns (GE Healthcare). All used oligonucleotides were in a double-stranded form, that was obtained using 1 µM of [^32^P]-labeled or 5 µM of unlabeled oligonucleotide solutions pre-incubated at 95°C for 15 min and then slowly cooled to 20°C.

**Table 1.**
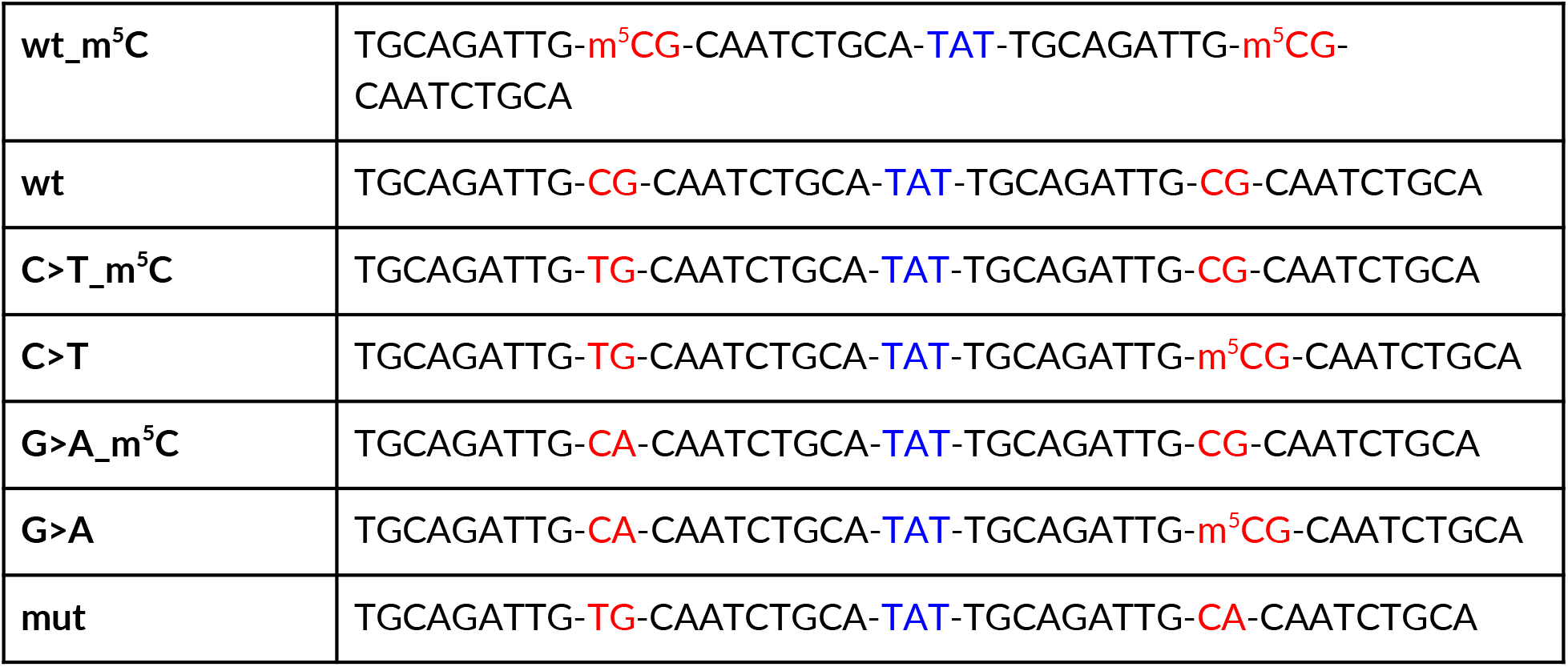
Oligonucleotides used for experimental verification of the CEBPB binding. The wild type (wt) consensus C/EBP binding site is palindromic, TAT serves as the loop. The core CpG (red) and the loop (blue) are between hyphens.

To avoid nonspecific binding of transcription factors, the nuclear extract was pre-incubated for 20 min at 30°C with the nonspecific Salmon Sperm DNA (Invitrogen): 1 µg of DNA per 1.5 µl of the nuclear extract (approximately 5 µg) in the reaction buffer (20 mM Hepes 7.6, 140 mM NaCl, 1.5 mM MgCl_2_, 1 mM DTT, 7% glycerol).

The reaction mixture contained [^32^P]-labeled wt_m^5^C and the appropriate preincubated nuclear extract (0.05 pmol [^32^P]-wt_m^5^C per 1.5 µl of original nuclear extract) in the reaction buffer. The mixture was incubated for 20 min at 30°C. The resultant DNA-protein complexes were separated in native 4% PAAG in 0.5x TBE (44.5 mM Tris, 44.5 mM boric acid, 1 mM EDTA) and visualized by autoradiography. The relative radioactivity was determined using a Packard Cyclone Storage Phosphor System (Packard Instrument Company, Inc.). For competition experiments, 2.5, 5, or 10 pmol (50, 100, or 200 fold molar excess) of unlabeled oligonucleotides was added simultaneously with [^32^P]-labeled wt_m^5^C to the reaction mixture. When necessary, 0.5 or 1 µg of anti-DDDDK tag antibodies was added to the nuclear extract before the preincubation step.

## Acknowledgments

This study was supported by the Russian Science Foundation grants 19-74-10079 (to I.A.E., wet-lab validation) and 17-74-10188 (to I.V.K., initial bioinformatics analysis), and Russian Foundation for Basic Research grant 18-34-20024 (to I.V.K., optimal motifs analysis). Structural analysis was performed under the Russian Ministry of Education and Science state project 0095-2019-0009. We thank E. Serebrova for the help in manuscript preparation.

## Author contributions

IVK and DP designed the study; IAE performed the wet-lab verification; ASE performed the mutation signature analysis; OSN performed the structural analysis; ADF performed initial cistrome analysis; IEV performed motif analysis; ASE, OSN, IEV, and IVK wrote the manuscript. All the authors read and approved the final manuscript.

## Expanded View Figure legends

**Figure EV1. Mutational signatures of stem cells.**

**A** Characteristics of the mutational signatures identified in the set of somatic mutations in hASCs and iPSC samples considering 96 context-dependent mutation types.

**B** Hierarchical clustering of samples based on contributions from individual mutation signatures.

**Figure EV2. Cancer mutations within regulatory regions are enriched within C/EBP motif occurrences.**

**A** Cosine similarity between hASC and iPSC mutation signatures and mutation profiles of selected cancer samples.

**B** Odds ratios (X-axis) demonstrating the enrichment of cancer mutations within C/EBP motif occurrences in the cistrome regions. Bar groups denote different cancer samples, the bars within each group denote mutations of signature S1 and [C>T]G mutations. The number of mutations within C/EBP occurrences in the cistrome is shown in brackets. Orange bars correspond to the P-values of less than 0.05 (Fisher’s exact test). Cancer samples: *COAD-US* - Colon Adenocarcinoma - TCGA, US; *READ-US* - Rectum Adenocarcinoma - TCGA, US; *STAD-US* - Gastric Adenocarcinoma - TCGA, US; *COCA-CN* - Colorectal Cancer - CN; *GACA-CN* - Gastric Cancer - CN.

