## Supplementary figures and images for "Enhanced C/EBPs binding to C>T mismatches facilitates fixation of CpG mutations"

### Fig EV1

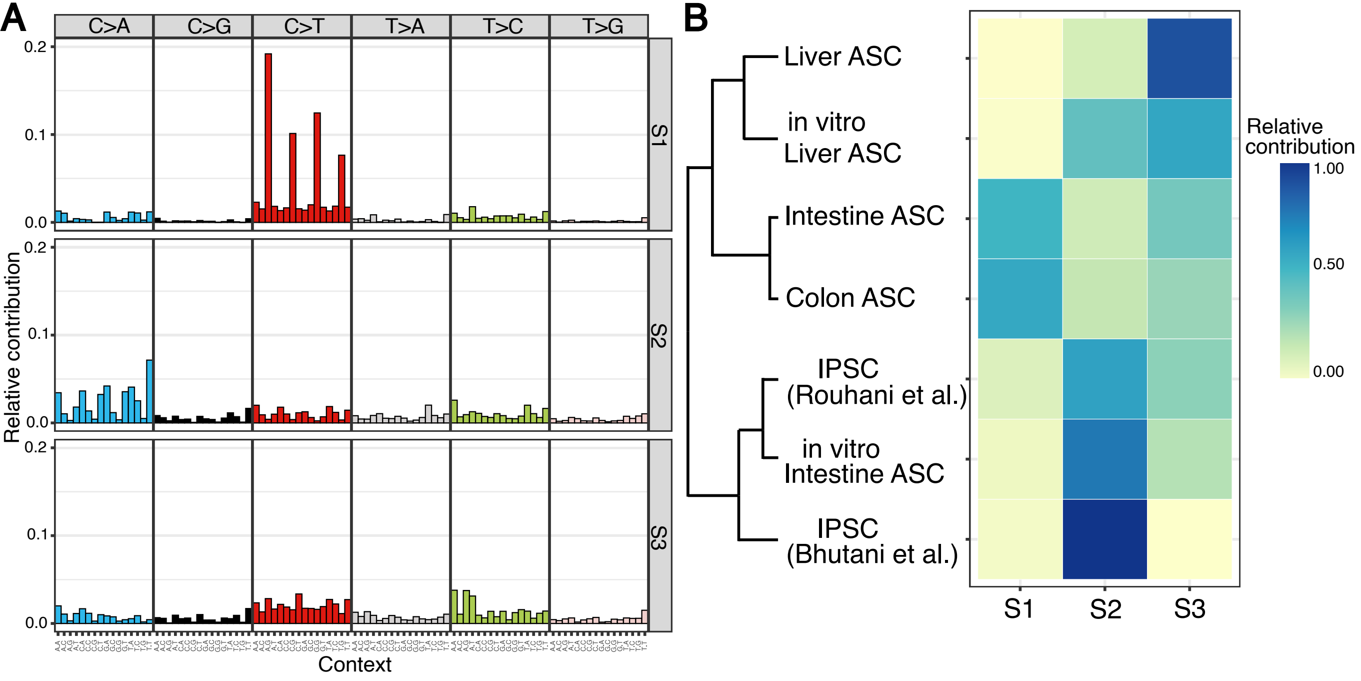

### Fig EV2

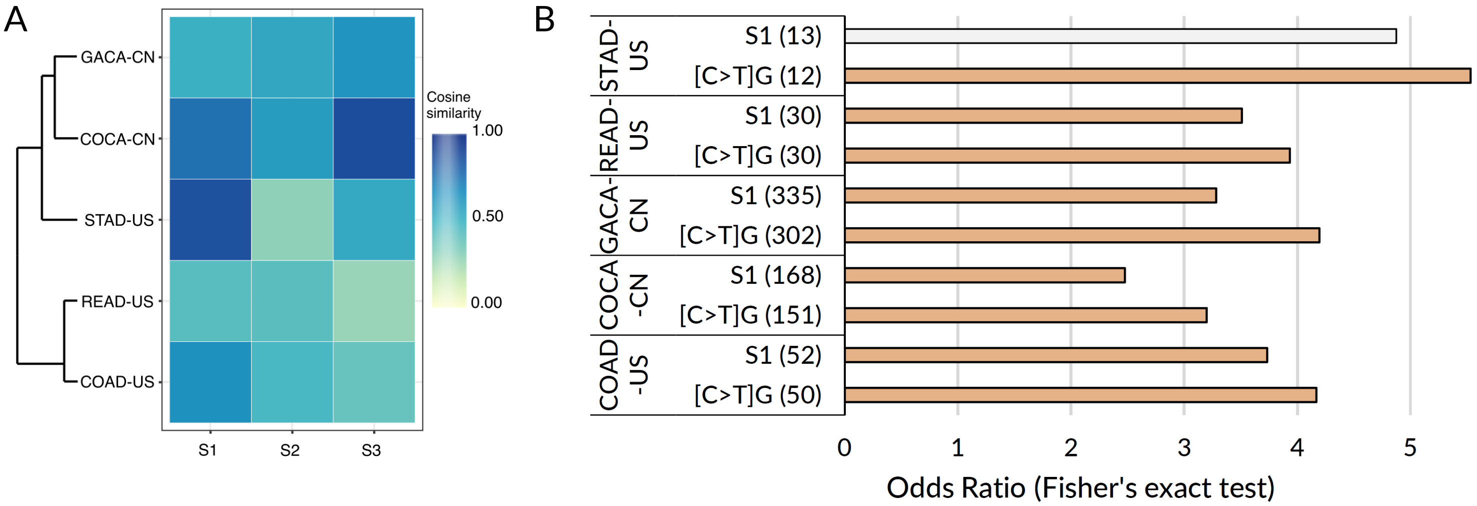
